# Larger fish upstream in a small stream: what are the causes of this longitudinal pattern?

**DOI:** 10.1101/2025.01.28.635151

**Authors:** Pedro Wolf, Piatã Marques, Rosana Mazzoni

## Abstract

Stream fishes often display distinct spatial patterns in size distribution, with one possible pattern being larger individuals dominating the headwaters. This “larger fish upstream pattern” (LFUP) has been widely documented, yet the ecological drivers remain unclear. We investigated factors contributing to LFUP in the fish community of a South American stream. Using historical data (1994–1998) from electrofishing surveys across six sites, we analyzed size distribution across eight species, with predictors including upstream distance, food availability, predation pressure, and intraspecific competition. Linear mixed models indicated that four species exhibit consistent LFUP, primarily driven by upstream distance or food availability. Species with pelagic or lithopelagic spawning displayed LFUP, supporting the hypothesis that LFUP is linked to reproductive strategies requiring upstream movement to maintain population stability. Our findings support that habitat connectivity and complexity are crucial for conserving small pelagic and lithopelagic fish species, as suggested in previous works. This study contributes to the field by highlighting mobile behavior in small tropical stream fishes and underscores the need for conservation strategies that account for movement patterns and habitat requirements in stream systems worldwide.

## Introduction

Stream fishes often show marked variation in body size distribution from the river mouth towards headwaters. A larger fish upstream pattern (LFUP) has been frequently described around the world (e.g. Jellyman & Graynoth, 1994; Hughes, 1999; Mazzoni et al., 2004; Perkin et al., 2023). In such cases, older, reproductively mature individuals often dominate upstream reaches, while juveniles are found at downstream sites (Hughes, 1999; Vitule et al., 2008). Multiple environmental factors can lead to patterns in fish size distribution (Harvey & Stewart, 1991; Giliam et al., 1993; Gurí et al., 2024). Disentangling such factors is important to expose specific drivers of the LFUP. Some hypotheses that attempt to expose specific drivers are Colonization Cycle Hypothesis (CCH) (Muller, 1982) and Static Population Growth and Mortality Gradients hypothesis (SPG) (Hughes, 1999).

The Colonization Cycle Hypothesis (CCH) (Muller, 1982) is a movement hypothesis evoked to explain the LFUP. According to this hypothesis, fish species whose eggs and/or larvae are vulnerable to downstream drift must compensate this by moving upstream to spawn. This behavior creates a gradient of larger individuals upstream, as they move to counteract the potential loss of offspring due to downstream drift. The CCH was initially developed for stream invertebrates and subsequently used to explain the size distribution of pelagic and lithopelagic spawning fish (Archdeacon et al., 2018; Platania et al., 2020; Steffensmeier et al., 2022; Perkin et al., 2023). These types of spawning involve floating eggs or free flowing larvae (Simon, 1999), that are subjected to significant downstream drift during their development (Platania & Altenbach, 1998; Platania, 2007). Therefore, this hypothesis considers the reproductive strategy and links reproduction to movement (Reynolds, 1983).

In addition to reproduction, other ecological factors can shape size distribution. The large fish upstream pattern can be a result of differential growth (Goto 1989). This mechanism is referred to as the Static Population Growth and Mortality Gradients hypothesis (SPG). This hypothesis posits that fish populations are stationary and longitudinal abiotic gradients (e.g. food availability, temperature) regulate growth (Hughes, 1999). Predation pressure and intraspecific competition can also play a role. The presence of predators regulates body size, by preying on specific size classes or as a life history response (Lundvall et al., 1999; Kristiansen et al., 2000; Gorini-Pacheco et al., 2017). Predation can also lead to larger prey size by reducing intraspecific competition and increasing resource availability per individual (Brönmark et al. 1995; Nakazawa et al., 2007).

In this paper we investigate the factors driving the LFUP pattern of the stream dwelling fish community. Our goal is to first assess wich species display LFUP. Then we will assess which ecological factor (distance upstream, differential growth, predation, intraspecific competition or reproductive strategy) better explain the observed pattern. Following the Colonization Cycle Hypothesis, we believe that the LFUP prior evidence for other species (e.g. Hughes, 1999; Mazzoni et al., 2004; Perkin et al., 2023) indicates larger fish moving upstream in our system. Thus, our hypothesis is that LFUP is linked to reproductive strategy of the fish. We expect that only species with pelagic and lithopelagic spawning will present a LFUP, and that distance upstream will be the primary predictor of it. Specifically.

## Methods

### Study area

We studied the size distribution off fishes in the Ubatiba stream (Figure 1). This stream is in the northeastern part of Rio de Janeiro State, in Brazil (22.8817° S, 42.8035° W). The fish diversity of the Ubatiba stream is composed of 24 species with different abundances across the stream system. For this study, we selected 8 species because of their wide distribution and abundance (Table 2). Fishes were sampled in 6 sites from downstream (site P6) to upstream (site P1). More details about the fluvial system can be found at Mazzoni & Lobon-Cerviá (2000) and Mazzoni et al. (2006).

**Figure 1.**
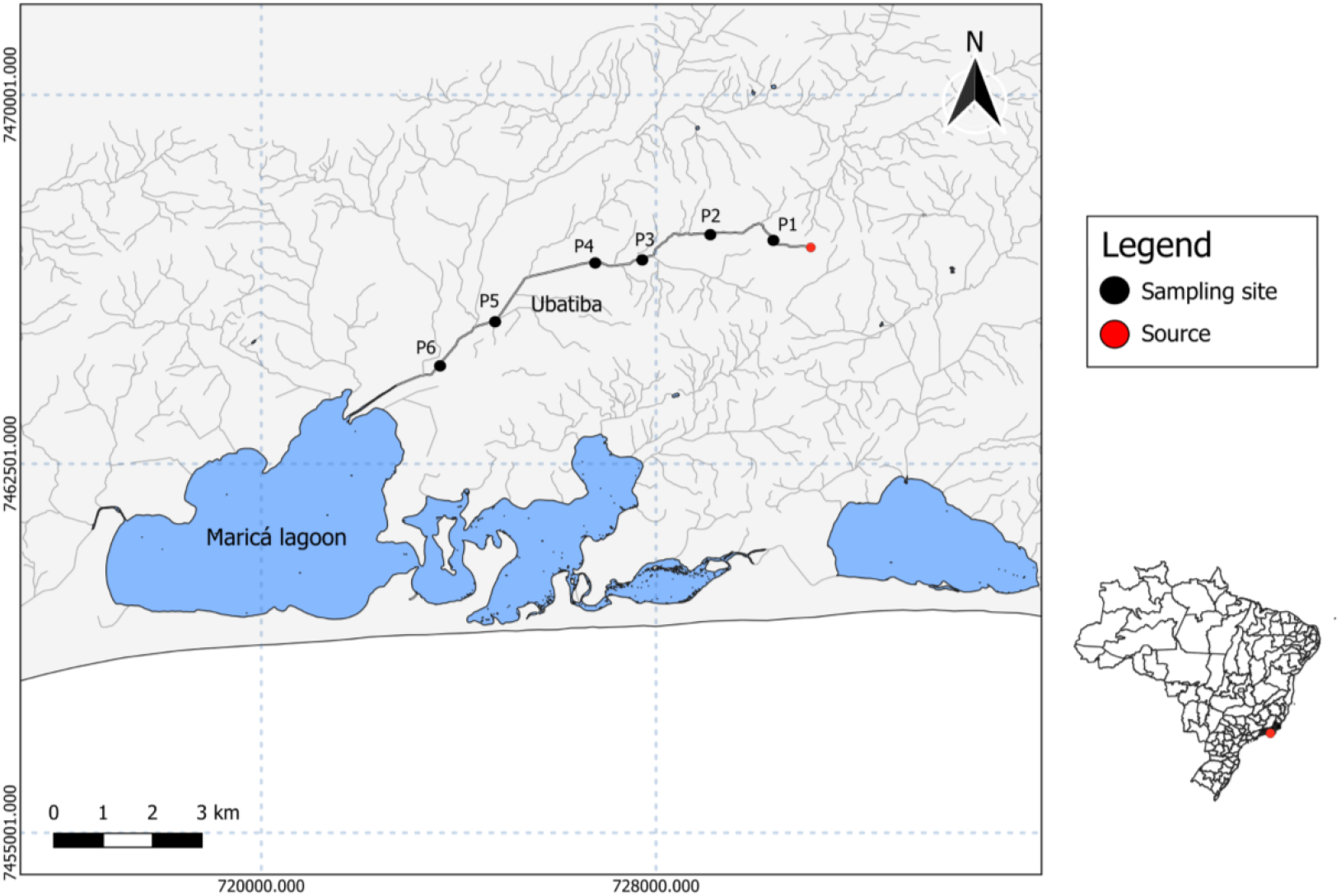
Ubatiba Fluvial System - Maricá (RJ). The map illustrates the Ubatiba River system along with the adjacent coastal lagoons. Sampling sites are arranged sequentially from downstream (P6) to upstream (P1). The stream source is highlighted in red, indicating that our farthest sampling site were positioned close to the headwaters.

### Field sampling techniques

The data for this study were derived from a historical dataset collected bimonthly between July 1994 and May 1998 across six sites in the Ubatiba system. Fish sampling was conducted through electrofishing (900 W, 220 V, 1-2 A) along ∼100 m stretches of the stream using the successive removal method (Zippin, 1958). Three successive removals were performed along each ∼100 m stretch, with the ends closed off using mesh nets (0.5 cm between nodes). Captured fish were held in floating cages to prevent recapture until the end of the process. Fish were identified to the species level, measured for standard length (SL, cm), and subsequently returned alive to the stream. A total of 33,145 fish from eight species were captured for this study. Information about the spawning types of these species was obtained from the literature (Table 1).

**Table 1.**
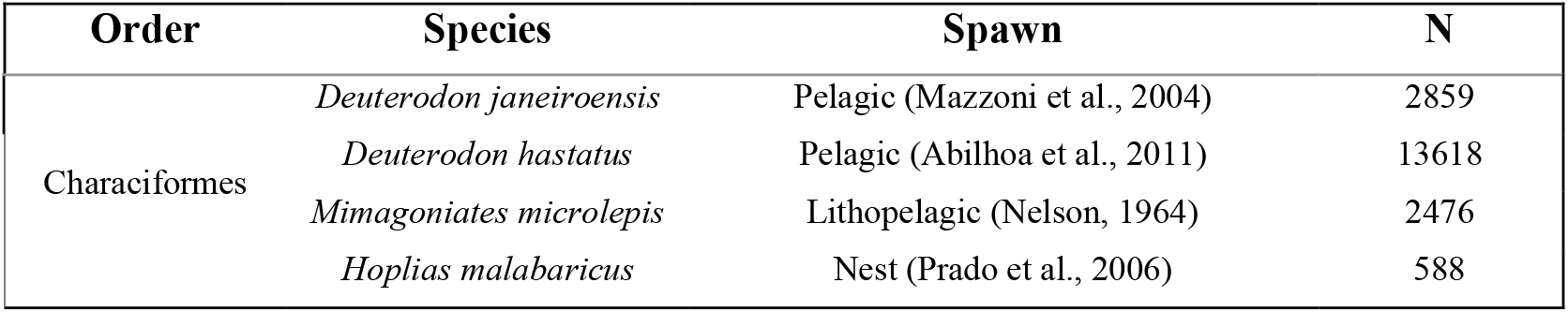
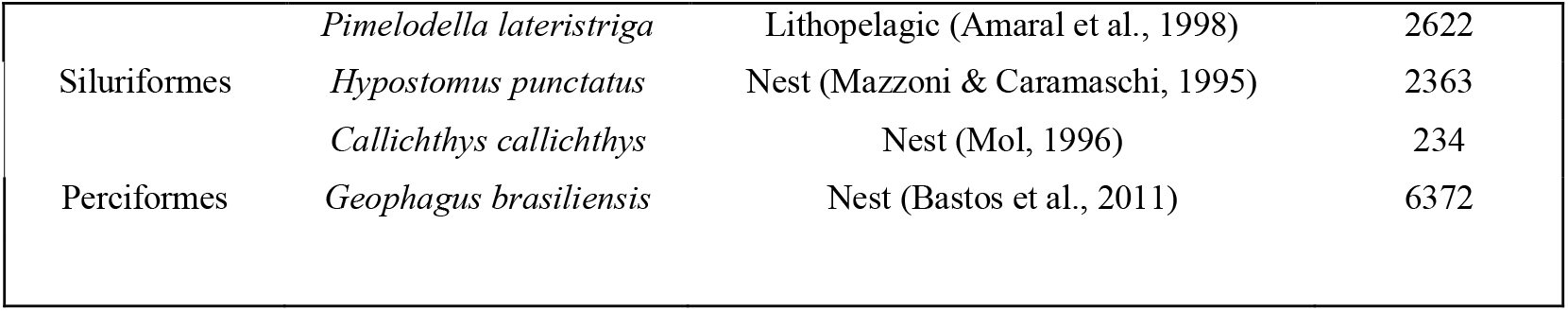
Fish species considered in this study, exposing the spawning strategy and their total abundances following the 5 years of study.

The distance of each site from the stream mouth was designated as “distance upstream” and measured using ARCGIS. The food availability on sites was parameterized with data extracted from Mazzoni & Lobón-Cervià (2000), accounting for the mean fish production rate (kg ha−1 yr−1) of each species at each site from 1994 to 1995. This information was used to rank sites according to their suitability for optimal species growth, from the least to the most favorable conditions. Predation pressure was assessed based on the abundance of the top piscivorous fish species *Hoplias malabaricus, Symbranchus marmoratus*, and *Rhamdia quellen*, which coexisted with the study species at each site and sampling occasion. Intraspecific competition was quantified as the abundance of coexisting conspecifics at the same site and sampling occasion. Although stream depth is regarded as a reliable predictor of fish size (Harvey & Stewart, 1991), it was excluded from the analysis, as the sites in Ubatiba have similar depths.

### Statistical analyses

First, we assessed which species show a larger fish upstream size distribution. For that, we built linear models using size (cm) as the response variable, and the distance upstream (km) as the predictor. The species without a significant size distribution upstream were removed from further analyses. In the second stage of analysis, we tested which predictor (i.e. distance upstream, food availability, predation or intraspecific competition) was the main driver of size distribution. For that, we built linear mixed models (LMM’s) (fixed slopes) for each species using size as the response variable, and distance upstream (km), production rate (kg ha^−1^ yr^−1^), number of predators and number of conspecifics as the fixed factors. Sampling date was included as random factor. We did not include predation as a factor when modeling *H. malabaricus* because this species is the main predator in the system. In addition, we built a neutral model using LMM to test if a neutral size better explain size distribution. The model includes size as the response variable, a neutral fixed factor (1) and the sampling date as the random factor. We compared each species model a neutral model using the Akaike information criterion (AIC). A significative difference between predictors and neutral models were considered when ΔAIC ≥ 2 (Burnham & Anderson, 2004).

We assessed the fit of each model by plotting residuals vs fitted values, and QQ-plots. The model’s family were selected based on data distribution. The LMMs were fitted using the lmer function of the lme4 package (Bates et al., 2024). The variance explained by both fixed plus random factors (conditional R square, R^2^c) and the variance explained only by fixed factors (marginal R square, R^2^m) were estimated using the r.squaredGLMM function of the “MuMIn” package for R (Barton, 2024). We also accessed the collinearity of the predictors using the function VIF from the “car” package for R (Fox & Weisberg 2010).

## Results

Our models showed that *Deuterodon janeiroensis, Deuterodon hastatus, Mimagoniates microlepis, Pimelodella lateristriga*, and *Hoplias malabaricus* have a significant Larger Fish Upstream pattern (Table 2). Conversely, *Geophagus brasiliensis* and *Hypostomus punctatus* exhibited the opposite trend, with larger individuals found near the stream mouth, and *Callichthys callichthys* showed no significant relationship between size and distance upstream.

**Table 2.** Results of the linear models used to test if fish size tend to increase in the longitudinal gradient. For each model, the only fixed factor was distance from the sampling site to the river mouth (Distance, km). Only species having larger fish upstream LFUP were used in further analysis (in bold).

| Species | Intercept | Estimate | F | p | R<sup>2</sup> |
| --- | --- | --- | --- | --- | --- |
| *Deuterodon janeiroensis* | <0.001 | 0.2 | 667 | <0.001 | 0.22 |
| *Deuterodon hastatus* | <0.001 | 0.1 | 4256 | <0.001 | 0.27 |
| *Mimagoniates microlepis* | <0.001 | 0.06 | 413 | <0.001 | 0.24 |
| *Hoplias malabaricus* | <0.001 | 0.16 | 8 | 0.005 | 0.01 |
| *Pimelodella lateristriga* | <0.001 | 0.06 | 124 | <0.001 | 0.05 |
| *Callichthys callichthys* | <0.001 | -0.02 | 0.2 | 0.63 | 0.004 |
| *Hypostomus punctatus* | <0.001 | -0.005 | 57 | <0.001 | 0.04 |
| *Geophagus brasiliensis* | <0.001 | -0.007 | 616 | <0.001 | 0.11 |

The second set of models was applied only to species showing a significant larger fish upstream pattern (Table 2). All models with predictors fitted better the data than neutral models (Table 4). Distance upstream was a significant predictor for all species, except *H. malabaricus* (Estimate = 0.04, SE = 0.06, F = 9, p = 0.6). For *D. janeiroensis* (Estimate = 0.15, SE = 0.009, F = 634, p < 0.001), *D. hastatus* (Estimate = 0.1, SE = 0.002, F = 4755, p < 0.001), and *P. lateristriga* (Estimate = 0.08, SE = 0.006, F = 266, p < 0.001), distance upstream was the primary predictor of size distribution (Table 3, Figure 2). Food availability was significant for all species (all p < 0.001) and was the primary predictor of larger sizes at sites for *M. microlepis* (Estimate = 0.13, SE = 0.02, F = 12, p < 0.001) and *H. malabaricus* (Estimate = 0.11, SE = 0.02, F = 27, p < 0.001) (Table 3). In *D. janeiroensis* (Estimate = 0.03, SE = 0.004, F = 16, p < 0.001) and *D. hastatus* (Estimate = 0.003, SE = 0.0006, F = 7, p < 0.001), food availability had a minor positive effect on length, while it had a slight negative effect on *P. lateristriga* (Estimate = -0.05, SE = 0.006, F = 73, p < 0.001). The sizes of *D. janeiroensis* (Estimate = 0.02, SE = 0.006, F = 5, p = 0.001) and *P. lateristriga* (Estimate = 0.01, SE = 0.003, F = 12, p < 0.001) tended to increase slightly at sites with higher predator abundance, while *M. microlepis* tended to be smaller in such sites (Estimate = -0.01, SE = 0.002, F = 12, p < 0.001). Finally, *D. janeiroensis* (Estimate = -0.01, SE = 0.001, F = 50, p < 0.001), *D. hastatus* (Estimat = -0.0004, SE = 0.00006, F = 46, p < 0.001), *M. microlepis* (Estimate = -0.003, SE = 0.006, F = 28, p < 0.001), *P. lateristriga* (Estimate = -0.001, SE = 0.0007, F = 4, p = 0.04), and *H. malabaricus* (Estimate = -0.12, SE = 0.05, F = 5, p = 0.02) tended to be smaller at sites with higher numbers of conspecifics.

**Table 3.** Results of the four LMM models used to assess the best predictors of body size (mm) for the different species. For each model, the fixed factors include the distance upstream (Distance, km), the sites food availability for species (Production, kg ha−1 yr−1), the number of predators (Predation, n), and the number of conspecifics (Intracomp), and sampling date as the random factor. It includes the marginal R^2^ (MR^2^), conditional R^2^ (CR^2^), and the effect, standard error (SE) p- and F-value for each predictor. The predictor with the greatest effect per model is highlighted in bold.

| Model | Predictors | Estimate | SE | F | p |
| --- | --- | --- | --- | --- | --- |
| *Deuterodon janeiroensis*<br>MR<sup>2</sup> = 0.21<br>CR<sup>2</sup> = 0.4 | Intercept | 3.9 | 0.2 |  | <0.001 |
|  | **Distance** | 0.15 | 0.009 | 634 | <0.001 |
|  | Food | 0.03 | 0.004 | 16 | <0.001 |
|  | Predation | 0.02 | 0.006 | 5 | 0.001 |
|  | Intracomp | -0.01 | 0.001 | 50 | <0.001 |
| *Deuterodon hastatus*<br>MR<sup>2</sup> = 0.29<br>CR<sup>2</sup> = 0.35 | Intercept | 2.64 | 0.06 |  | <0.001 |
|  | **Distance** | 0.10 | 0.002 | 4755 | <0.001 |
|  | Food | 0.003 | 0.0006 | 7 | <0.001 |
|  | Predation | -0.001 | 0.001 | 3 | 0.3 |
|  | Intracomp | -0.0004 | 0.00006 | 46 | <0.001 |
| *Mimagoniates microlepis*<br>MR<sup>2</sup> = 0.33<br>CR<sup>2</sup> = 0.42 | Intercept | 2.59 | 0.07 |  | <0.001 |
|  | Distance | 0.04 | 0.005 | 405 | <0.001 |
|  | **Food** | 0.13 | 0.02 | 12 | <0.001 |
|  | Predation | -0.01 | 0.002 | 27 | <0.001 |
|  | Intracomp | -0.003 | 0.0006 | 28 | <0.001 |
| *Pimelodella lateristriga*<br>MR<sup>2</sup> = 0.12<br>CR<sup>2</sup> = 0.32 | Intercept | 5.89 | 0.17 |  | <0.001 |
|  | **Distance** | 0.08 | 0.006 | 280 | <0.001 |
|  | Food | -0.06 | 0.006 | 82 | <0.001 |
|  | Predation | 0.01 | 0.003 | 10 | <0.001 |
|  | Intracomp | -0.001 | 0.0007 | 4 | 0.03 |
| *Hoplias malabaricus* | Intercept | 11.4 | 0.8 |  | <0.001 |
| MR<sup>2</sup> = 0.07 | Distance | 0.04 | 0.06 | 9 | 0.6 |
| CR<sup>2</sup> = 0.12 | **Food** | 0.11 | 0.02 | 27 | **<0.001** |
|  | Intracomp | -0.12 | 0.05 | 5 | **0.02** |

**Table 4.** Model comparison of the six mixed effect models (LMM) by the AIC method and adjusted R^2^ of the predictor’s models. The ΔAIC (≥ 2) are highlighted in bold.

| Species | Model | AIC | ΔAIC |
| --- | --- | --- | --- |
| *Deuterodon janeiroensis* | Neutral | 9844 |  |
|  | Predictors | 9269 | **575** |
| *Deuterodon hastatus* | Neutral | 27834 |  |
|  | Predictors | 23880 | **3954** |
| *Mimagoniates microlepis* | Neutral | 2534 |  |
|  | Predictors | 2177 | **357** |
| *Pimelodella lateristriga* | Neutral | 8058 |  |
|  | Predictors | 7755 | **303** |
| *Hoplias malabaricus* | Neutral | 3315 |  |
|  | Predictors | 3295 | **20** |

**Figure 2.**
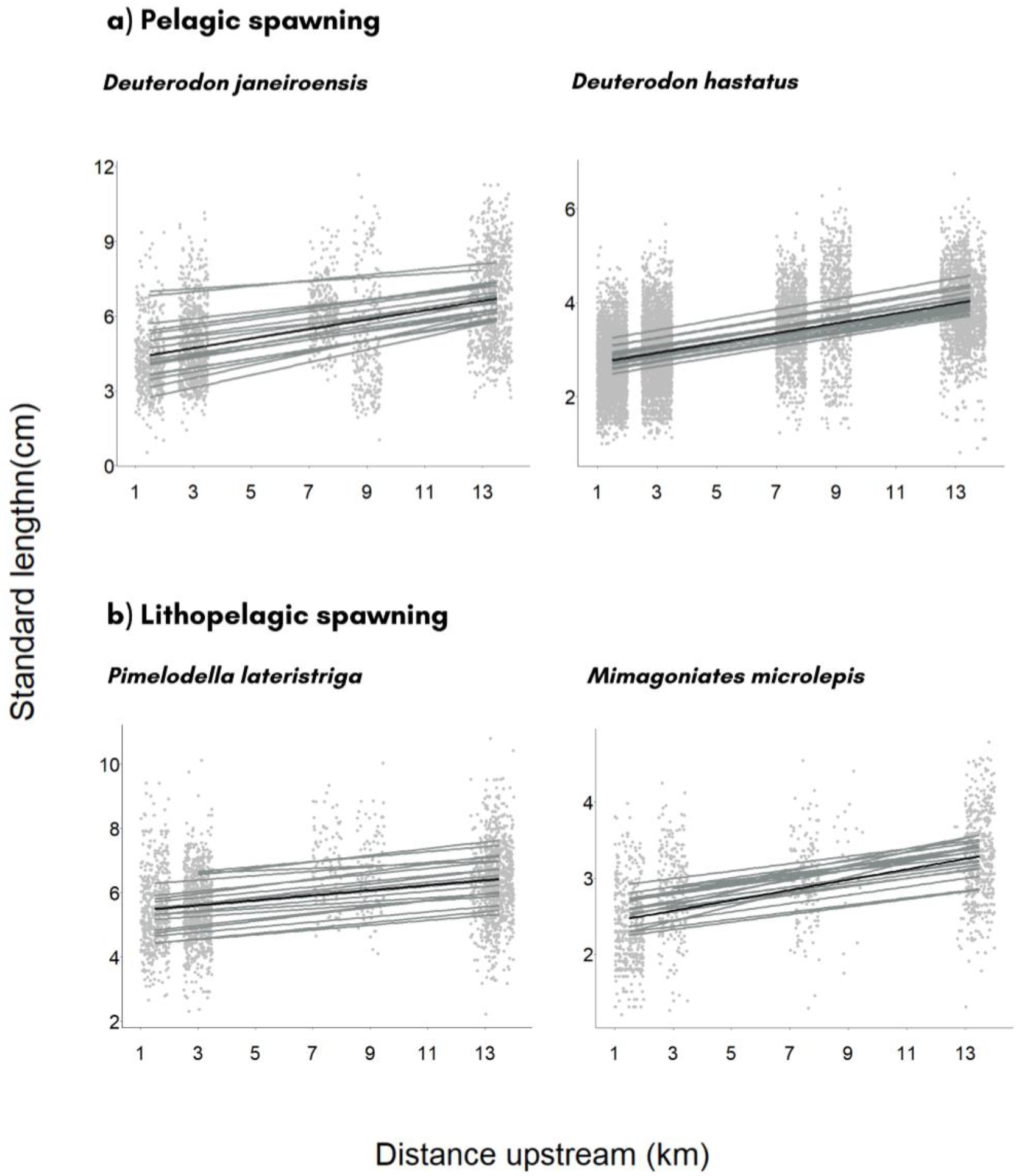
Marginal fits (black lines) from the mixed effect models with fixed intercepts, and the conditional fits (grey lines) for each sampling campaign.

## Discussion

Our results indicate that most species align with our expectations that only pelagic and lithopelagic fish present a LFUP pattern. Both *Deuterodon* species (pelagic spawners) and *Pimelodella lateristriga* (lithopelagic spawner) exhibit a larger fish upstream pattern (LFUP), which is primarily driven by upstream distance compared to other ecological factors. The only exception was *Mimagoniates microlepis*, a lithopelagic spawner, which deviated from our prediction presenting food availability as the primary predictor. However, there are plausible explanations for this deviation, and further investigation is necessary. Finally, also as expected, species exhibiting some levels of parental care do not display LFUP.

Recolonization of upstream segments of river and streams is presumed necessary for the continued occupancy of upstream reaches in species vulnerable to downstream displacement (Wilde & Urbanczyk, 2014; Steffensmeier et al., 2022; Perkin et al., 2023; Steffensmeier et al., 2024). A colonization cycle of upstream areas seems to be the strategy these species display to compensate offspring downstream displacement and maintain the population in upstream sections (Muller, 1982). The LFUP is a size distribution that reflects this process through space and time (Abilhoa et al., 2011). Prior movement research in Ubatiba fish community shows that pelagic and lithopelagic species are highly mobile, with *Deuterodon* species displaying upstream biased movement and *P. lateristriga* augmented non-directional movement (Mazzoni et al., 2012; Mazzoni et al., 2018). First it was believed that upstream biased movement was essential for pelagic and lithopelagic fish (Moore, 1944). However, our findings add to the growing evidence that recolonization by only a subset of individuals, caused by augmented non-directional movement, is sufficient to sustain the population in upstream sections (Speirs & Gurney, 2001; Naman et al., 2016; Steffensmeier et al., 2022; Steffensmeier et al., 2024).

We expected *M. microlepis* to show evidence of movement because it is a lithopelagic species. This can be a case of drift paradox., a concept created by Hershey et al. (1993) to refer to cases when species vulnerable to downstream displacement lack evidence of upstream recolonization. However, *M. microlepis* has a particular behavior of laying eggs in the abaxial face of instream vegetation (Nelson, 1964). One of the proposed solutions for the drift paradox is the partial retention of eggs and larvae in the stream (Speirs & Gurney, 2001; Naman et al., 2016). This can solve the drift paradox for *M. microlepis*, because the active laying of eggs in instream vegetation, a retentive structure responsible to reduce flow impact, it’s probably an adaptation to avoid downstream drift (Tabacchi et al., 2000). Hence, the patterns we found for *M. microlepis* reinforces that considering species specific traits is essential when evaluating the necessity of high mobile behavior in pelagic and lithopelagic species.

The three models of *D. janeiroensis, D. hastatus* and *P. lateristriga* had low marginal explanation power (MR^2^ = 0.21, 0.29 and 0.12, respectively). This outcome may have multiple explanations. Floods can drag large mid-water fish downstream (Hakamada & Penha, 2014; Smith & Kwak, 2015), what may coincide with the moment of sampling since lots of them were made after raining periods. Also, upstream movement in our case and in other works is not hypothesized to be caused by lots of synchronized individuals, therefore we could have caught individuals in downstream sections during upstream moving process. In addition, smaller individuals can be found upstream due to egg or larvae retention in pools, instream vegetation and rock crevices. Thus, pelagic and lithopelagic fish size distribution is a dynamic process that express itself in terms of proportion upstream versus downstream (Abilhoa et al., 2011), with larger individuals permanently dealing with the flow dynamics and younger forms drifting or being retained upstream.

The remaining species (*Callichthys callichthys, Hoplias malabaricus, Hypostomus punctatus* and *Geophagus brasiliensis*) likely avoid the downstream displacement of individuals by other mechanisms than movement. First, they can avoid the downstream displacement of offspring by nesting in slow-flow habitats and actively guarding their eggs and newborns (Mazzoni & Caramaschi, 1995; Mol, 1996; Prado et al., 2006; Bastos et al., 2011). This strategy aligns with the traditional time/energy allocation hypothesis (Bonte et al., 2006; Comte & Olden, 2018), which posits that investing energy into certain activities (e.g. parental care) reduces the energy available for other behaviors (e.g. reproductive movement). Second, adults of this species are benthonic species with behavioral (boundary layer use or lair establishment) and morphological (deep body shape) adaptations to deal with high flow velocities (Carlson & Lauder, 2011; Meyers & Belk, 2014), which make then resistant to floods (Webbs et al. 1996; Tew et al., 2002; Carlson and Lauder 2010). Thus, these species are not required to be highly mobile to deal with high flow.

Our results have important implications for the conservation of small pelagic and lithopelagic species, since high mobile behavior seems to be an important species trait. High mobile fish species are one the most threatened organisms on the planet according to the WWF Living Planet Index for the group (Deinet et al., 2020), and pelagophil species are the most imperiled among the ichthyofauna, especially when they have upstream biased movement (Steffensmeier et al., 2024). Major threats for this species are (1) loss of habitat complexity, which tends to remove upstream retentive structures for eggs and larvae (Worthington et al., 2014), and (2) fragmentation of river or stream connectivity, which hinders movement behavior and may be responsible for species demise from certain stream sections (Platania & Altenbach, 1998). Therefore, for improved management and conservation of pelagic and lithopelagic species, future research should build on previous findings, which suggest two key areas of focus: (1) understanding how habitat complexity supports recruitment in upstream areas and benefits fish populations (Medley & Shirley 2013; Valdez et al. 2019, 2021); and (2) enhancing fish movement tracking to determine the necessary longitudinal span to guide conservation actions (Dudley & Platania, 2007; Wilde & Urbanczyk, 2013; Allen & Singh, 2016; Steffensmeier et al., 2024). This is especially urgent in tropical streams, where there’s lack of information in fish movement ecology (Rasmussem & Belk, 2017; Comte & Olden, 2018; Mazzoni & Barros, 2021).

